# Excessive confidence in hypomania arises from mood-biased reward integration, not a global shift in reward expectancy

**DOI:** 10.1101/2024.11.18.624111

**Authors:** Liam Mason, Sascha P. Woelk, Eran Eldar, Robb B. Rutledge

## Abstract

Mood disturbances are accompanied by marked and apparently non-rational changes in self-efficacy and goal-directed behaviour, yet the mechanisms linking affective states to these fluctuations remain unclear. Here we tested whether mood biases confidence in reward-based decision-making and whether this is amplified by trait instability in mood and self-efficacy. By formalising both confidence and mood within a computational model of reward learning, we adjudicated between two accounts of these clinically relevant effects: mood imposing a direct, evidence-independent shift in confidence, or mood distorting the accumulation of reward learning signals from which confidence is then rationally inferred.

Across two independent studies—a laboratory experiment (n = 35) and a preregistered online replication (n = 106)—participants completed a reinforcement-learning task with mood manipulation and repeated confidence ratings. Mood systematically biased confidence without affecting objective accuracy. Critically, this effect accumulated gradually across learning rather than emerging immediately after mood induction, even as mood itself returned toward baseline. These dynamics were captured by a computational model in which mood biases the accumulation of reward probability estimates. A single mood-bias parameter accounted for shifts in confidence, biased valuation of stimuli learned under different mood states, and amplification of these effects in individuals with elevated hypomanic traits.

These findings suggest that disturbances in confidence arise from mood-biased learning signals, providing a mechanistic account linking these distortions to disrupted goal pursuit and self-efficacy in mood disorders.

## Introduction

Intuitively, moods shape not only the decisions we make, but how confident we feel in taking them. Clinically, elevated mood is often accompanied by overconfidence and perceived ability to attain goals, while low mood is marked by doubt and reduced goal pursuit^1^. Drawing on evidence that mood and reward learning processes recursively influence each other^2–4^, we reasoned that momentary confidence—conceptualised here as an expression of self-efficacy during learning—would be impacted by these neurocomputational mechanisms.

Prior work demonstrates a bidirectional relationship between action outcomes and mood, where mood both reflects the cumulative impact of outcomes and feeds back to influence how subsequent outcomes are perceived. Specifically, mood appears to bias the integration of rewards, modulating the valuation and learning from outcomes^2–4^. Functional neuroimaging studies show that this “mood bias” operates at the level of striatal reward prediction errors (RPEs)^5–7^. This mood-contingent amplification of striatal RPEs is stronger as a function of hypomanic personality traits^5^, an established psychometric risk factor for bipolar disorder^8,9^ that is characterised by marked shifts in mood^9^, reward sensitivity^10–13^, and confidence^14^. In a recent model-based fMRI study, we confirmed that this “mood bias” mechanism is exaggerated in bipolar disorder patients out-of-episode, where momentary increases in positive mood amplified striatal responses to positive RPEs and periods of negative mood amplified responses to negative RPEs, compared to controls^7^.

These recursive dynamics, where mood and learning mutually reinforce one another, extend prior evidence of trait reward hypersensitivity in bipolar disorder^15–17^, and are consistent with our theoretical account of mood instability^18^. An important but untested corollary of this account concerns self-efficacy, the belief in one’s ability to produce desired outcomes, which is frequently perturbed in mood disorders, manifesting as grandiosity during manic states and hopelessness or self-doubt during depressive episodes^1^. If mood biases how reward outcomes are integrated, then mood-contingent distortions in learning signals are a plausible mechanism through which instability in mood and self-efficacy co-emerge.

One way to formalise mood-dependent fluctuations in self-efficacy is through confidence. Considerable work has shown that evolving value and action utility are reflected in confidence judgements^19–22^. The relationship with decision-making processes is bidirectional, where confidence has been shown to amplify confirmatory evidence and discount disconfirmatory information in both perceptual decision-making^23^ and reinforcement learning^24^. Importantly, there is evidence that confidence in these contexts may be subject to affective influences, at least in perceptual decision-making contexts. Confidence about motion signals was sensitive to arousal (disgust cues)^25^ and was also modulated by mood symptoms, despite intact performance accuracy, indicating a selective effect on confidence processes rather than task competence^22^. This dissociation is theoretically significant: recent frameworks distinguish between ‘local’ confidence in individual decisions and ‘global’ self-performance estimates reflecting broader beliefs about skills and abilities^26^, raising the possibility that mood selectively disrupts specific levels of this hierarchy. At the neural level, recent fMRI studies show that ventromedial prefrontal cortex (VMPFC) activity during decision-making tracks subjective confidence more closely than objective reward value acquired through reinforcement learning^27^. Positive affect has been shown to increase success expectancy in individuals with elevated hypomanic traits^14^. These studies motivate the present study’s aim of elucidating the computational mechanisms underlying the link between mood and confidence.

Here we tested a mechanistic account of how mood shapes momentary confidence during learning. Building on evidence that mood biases the integration of reward outcomes^2^,^5^, we asked whether mood influences confidence indirectly, by distorting how reward probabilities are accumulated over time, rather than imposing a global, evidence-independent shift in certainty. Under a learning-mediated account, mood-biased confidence should be reflected not as an immediate shift in certainty, but as a divergence in the trajectory of confidence over learning, emerging gradually across trials as value estimates are accumulated. Moreover, if this process reflects a vulnerability to affective modulation of inference, these confidence trajectories should be amplified as a function of hypomanic traits, consistent with heightened susceptibility to mood-linked distortions in self-efficacy.

To evaluate this account, we formalised a computational model in which mood, reward probability learning, and confidence are integrated within a single framework, and contrasted it with models in which mood acts directly on confidence. Grounded in prior work demonstrating mood-dependent biases in reward integration^2^,^5^, this approach allowed us to test whether observed confidence dynamics—and their modulation by trait instability in mood and self-efficacy^9,14^ —are better explained by mood-biased accumulation of reward evidence than by a global shift in certainty. Across a laboratory experiment and a preregistered online replication, we find converging evidence that transient mood states distort confidence through biased accumulation of reward likelihood rather than through a global shift in reward expectancy, and that this bias is amplified in individuals with elevated hypomanic traits.

## Results

### Confirmation of learning and mood manipulation

In both studies, participants completed a reinforcement learning task with mood manipulation (Figure 1). Planned analyses (see Supplementary Materials for full statistics) confirmed that participants chose the higher-probability option significantly more often than chance (all p < 10^−5^), and that positive and negative mood manipulations produced significant and opposing changes in momentary happiness in both samples (all p ≤ 0.0075).

**Figure 1.**
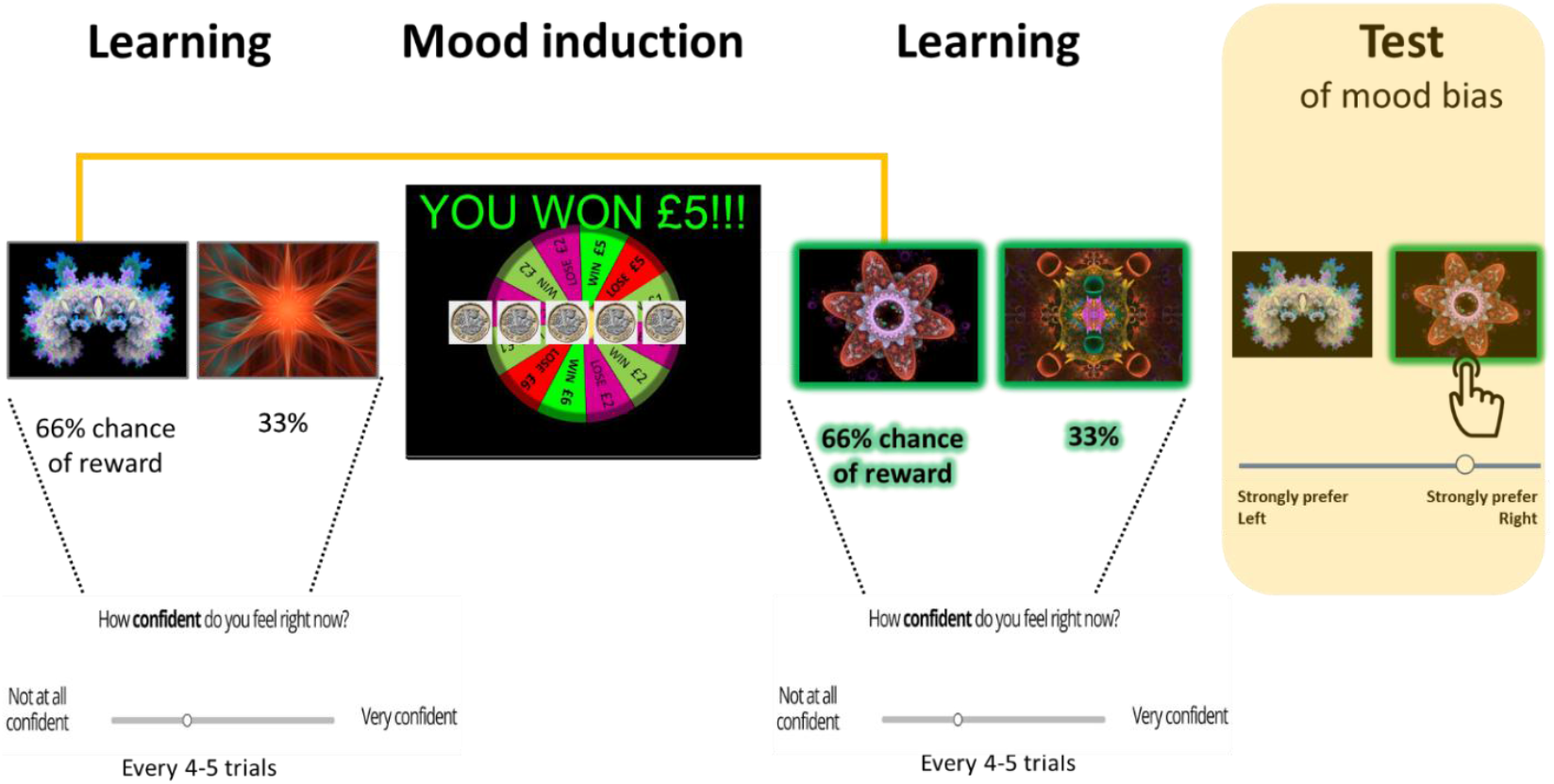
Experimental design. Participants completed two blocks of learning, one pre- and one post-mood induction. In each block, participants chose between two options which yielded low (33%) or high (66%) probability of reward. We tested whether mood induction biased reward learning of the subsequently encountered stimuli (green outline, not visible to participants, represents the predicted effect of a positive mood induction increasing valuation of the block 2 stimuli and biasing their preference in the between-block pairwise choices in the test block). Participants gave periodic confidence ratings after feedback (every 4-5 trials, pseudo-randomly, 10 ratings per learning block) where they were instructed to give their level of confidence about which option was superior.

### Mood biases momentary confidence, amplified by trait instability in mood and self-efficacy

There were no differences in confidence between groups prior to the mood manipulation [Study 1: t(33) = 0.63, p = 0.54; Replication study: t(103) = 0.047, p = 0.96]. Importantly, we also confirmed that there was no difference in objective performance between participants who experienced positive or negative mood manipulations in Study 1 [block 1: t(33) = 0.49, p = 0.63; block 2: t(33) = 0.41, p = 0.68], or in Replication study [block 1: t(103) = 1.11, p = 0.27; block 2: t(103) = 0.65, p = 0.51]. The results of the linear mixed effects model (Table 1) confirmed that confidence increased with learning (cumulative proportion of correct choices made). Confirming Hypothesis 1a, mood block exerted an effect on confidence, independently of objective trial-wise accuracy (Study 1: p=0.003, Replication: p=0.001). In both experiments, increases in confidence were observed under positive mood manipulation and decreases under negative mood manipulation (Figure 2, left column). Hypomanic traits, which we used to index trait instability in mood and self-efficacy (see Methods), showed a significant interaction with mood manipulation in both studies (Study 1: p=0.003; Replication: p=0.023). Confirming Hypothesis 1b, the effect of mood on confidence scaled with hypomanic traits (Figure 2, right). Moreover, the three-way interaction (learning × mood manipulation × hypomanic traits; p=0.039) shows that the stronger effect of mood in participants with elevated hypomanic traits increased over the course of learning (Supplementary Figure 1).

**Table 1.** Linear mixed effects modelling of the influence of mood manipulation in predicting subjective confidence in Study 1 and Replication study, accounting for trial-wise objective accuracy and hypomanic traits.

| Predictor | Study 1 |  |  |  | Replication study |  |  |  |
| --- | --- | --- | --- | --- | --- | --- | --- | --- |
|  | Beta | SE | *t* | *p* | Beta | SE | *t* | *p* |
| Intercept | -0.083 | 0.065 | -1.291 | 0.197 | 0.160 | 0.090 | 1.789 | 0.074 |
| Accuracy<sub>t</sub> | 0.692 | 0.117 | 5.918 | **<10<sup>-9</sup>** | 0.232 | 0.057 | 4.091 | **<10<sup>-5</sup>** |
| Hypomania | 0.000 | 0.000 | 0.172 | 0.863 | 0.004 | 0.003 | 1.382 | 0.167 |
| Moodblock | 0.253 | 0.086 | 2.946 | **0.003** | 0.0597 | 0.025 | 2.359 | **0.001** |
| Accuracy<sub>t</sub> × Moodblock | 0.480 | 0.574 | 0.837 | 0.403 | 0.396 | 0.210 | 1.891 | 0.059 |
| Moodblock × Hypomania | 0.001 | 0.000 | 3.012 | **0.003** | 0.008 | 0.003 | 2.272 | **0.023** |
| Accuracy<sub>t</sub> × Moodblock × Hypomania | -0.001 | 0.001 | -0.595 | 0.552 | -0.014 | 0.007 | -2.071 | **0.039** |

**Figure 2.**
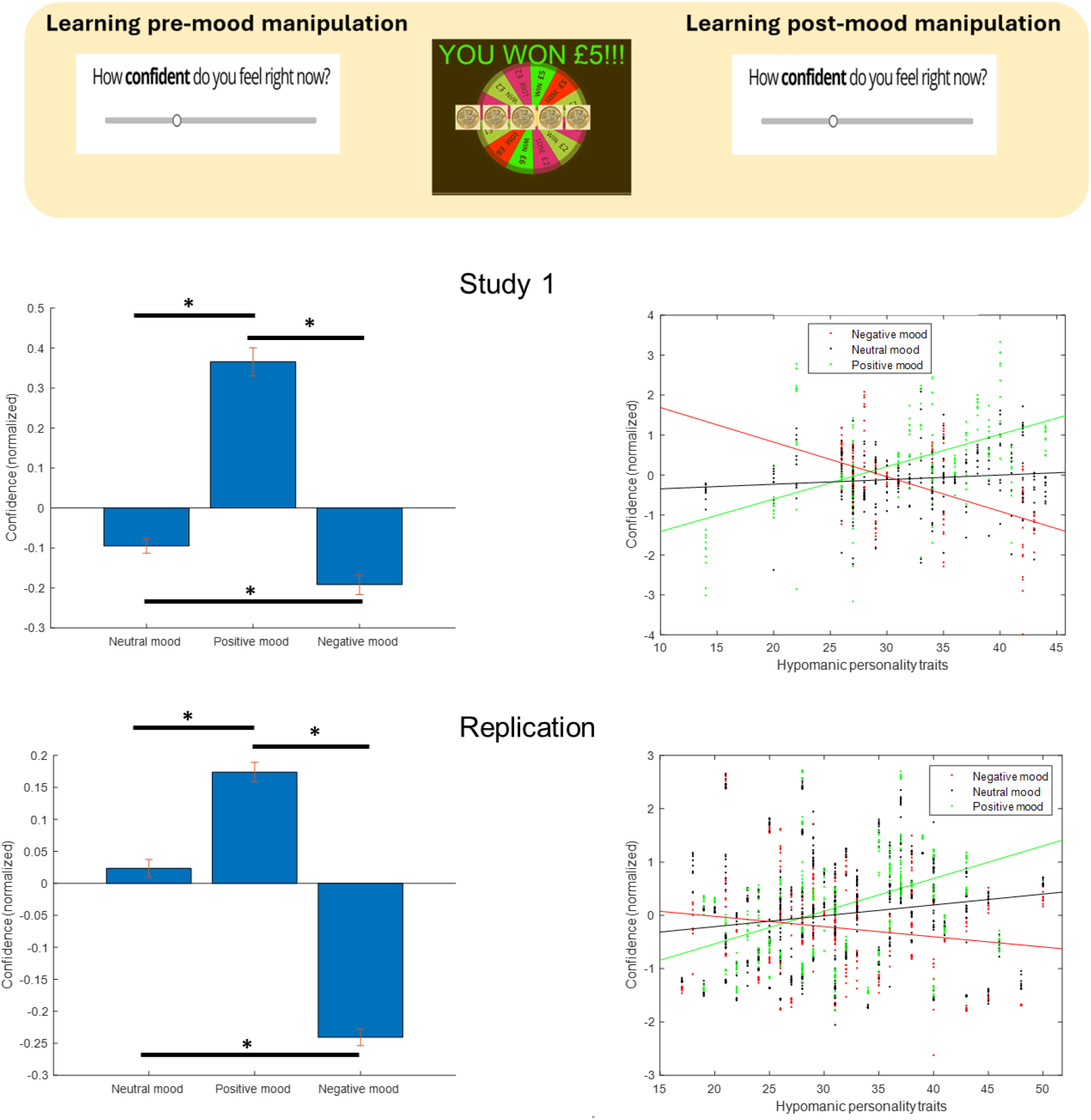
Mood biases subjective confidence across two experiments. **Left:** Relative to neutral mood, ie prior to mood manipulation, momentary confidence was increased under positive compared to negative mood manipulation, despite no differences in objective accuracy. Top: Study 1, Bottom: Replication study; error bars denote ± SEM. **Right:** Participants with higher hypomanic traits were more susceptible to mood biasing their confidence. *Note*: Predicted confidence is plotted from linear mixed effects model where all regressors are normalized, but raw HPS scores are shown for interpretability. Each point represents an individual confidence rating obtained at intermittent prompts during the task, with multiple ratings contributed by each participant.

### Parsing momentary confidence from momentary happiness

Because confidence ratings were collected post-outcome, they could in principle reflect transient affective responses to reward rather than a distinct inferential signal. To address this possibility, we re-estimated the above regression model while additionally accounting for the outcome on the preceding trial. As expected, outcome was indeed an additional predictor of confidence (in both experiments; *p* ≤ 0.0003) but, importantly, all prior mood and hypomanic trait effects remained (*p* ≤ 0.02; Supplementary Table 2), ruling out that these effects were primarily driven by affective responses to outcomes.

We next sought to establish that momentary confidence represents a distinct signal to momentary happiness. The influence of mood on confidence emerged progressively over the learning block, peaking late in learning (Figure 3b, right), in contrast to happiness, which showed its largest change immediately after mood induction and decayed thereafter (Figure 3b, left). This dissociation indicates that the mood manipulation shaped confidence through its interaction with learning, consistent with simulations of the Moody Accumulation model (Figure 3c).

**Figure 3.**
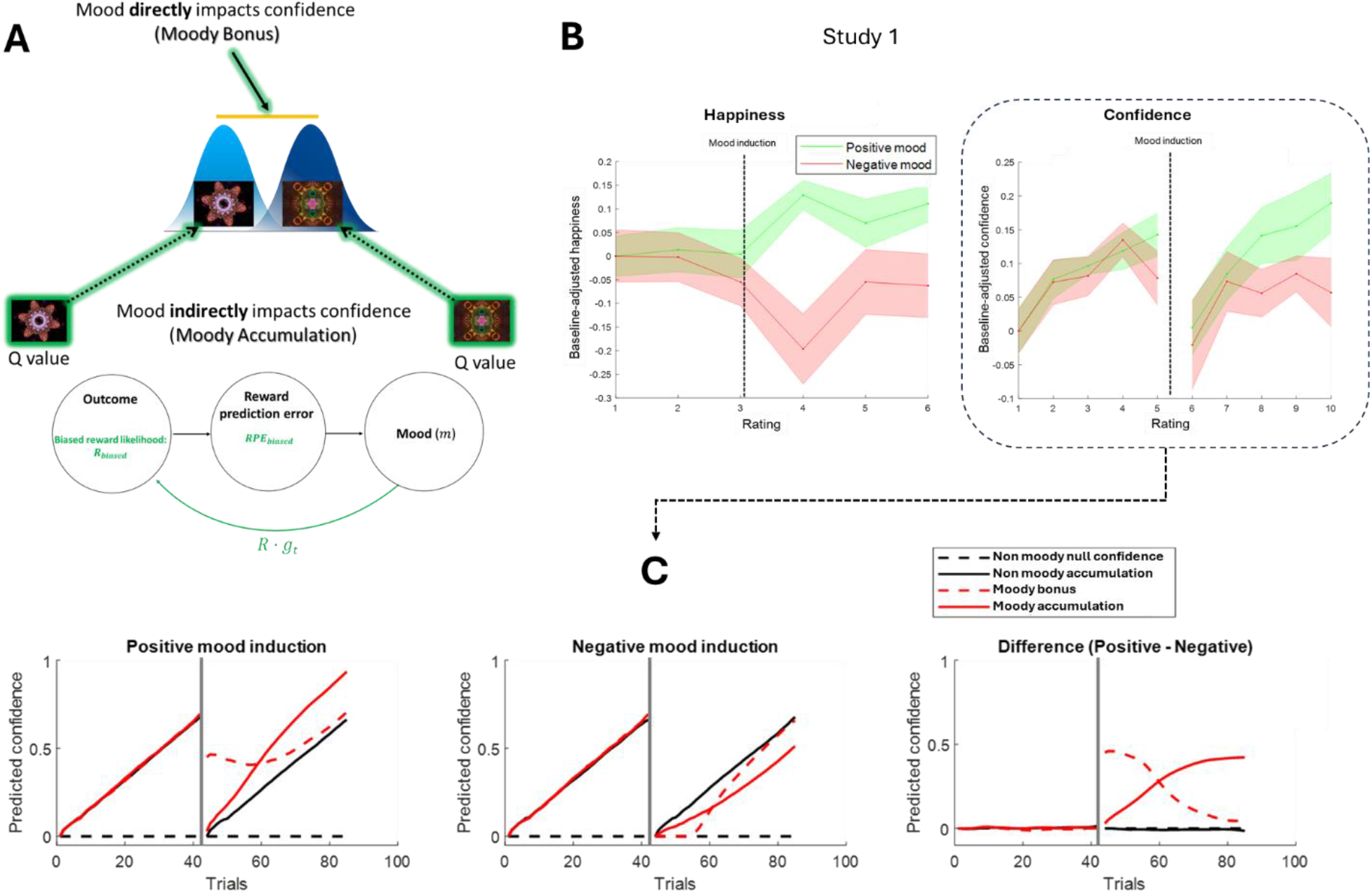
Computational simulations of mood-biased learning reproduce target empirical confidence dynamics. **A) Schematic depicting two candidate computational models for the impact of mood on momentary confidence**. In both models, momentary confidence is formalised as the trial-by-trial difference between the estimated reward probabilities of the available options (horizontal orange line); confidence is greater when these probability estimates become further apart through learning. The two models differ in where mood exerts its influence: in Moody Bonus (top), mood acts as a direct additive shift on confidence, on top of what is determined by reward learning. Conversely, in Moody Accumulation (bottom), mood biases the reward probability estimates themselves during learning, shaping confidence indirectly, through mood-distorted evidence accumulation. **B) Empirical data demonstrating that mood influences the accumulation of confidence**. Whilst the mood manipulation had its peak effect immediately post-induction and decayed back to baseline (left), its effect on confidence followed the opposite time course, reaching a peak at the end of the learning block (right). **C) Model simulations showing momentary confidence across two learning sessions, separated by mood induction** (a large reward prediction error at the dashed vertical line). Confidence increases throughout each session as reward probabilities are learned. Importantly, the two candidate models make opposing predictions about the temporal dynamics of mood’s influence on confidence. In the Moody Bonus model, mood has an immediate effect on confidence proximal to induction, whereas in the Moody Accumulation model, the effect emerges gradually in concert with learning, recapitulating the empirical time course shown in B.

Finally, we directly tested whether confidence and happiness index distinct constructs by sampling happiness at the same frequency as confidence (10 ratings per block) in a post-experiment learning block with novel stimuli. These measures showed clearly dissociable dynamics. First, whereas confidence increased over the course of the control block, mirroring the pattern observed in the learning blocks of the main experiment, happiness did not increase during the control block (Supplementary Figure 2). Second, participants’ happiness ratings in the control block were unrelated to their confidence ratings in the main task blocks (block 1: *r* = 0.10, *p* = 0.78; block 2: *r* = 0.02, *p* = 0.96), whereas confidence ratings were strongly correlated across the two main task blocks (*r* = 0.74, *p* = 0.02). Together, these findings indicate that momentary confidence is dissociable from momentary happiness, consistent with our claim that confidence tracks a learning-dependent inferential signal shaped by mood.

### Mood during learning biases subsequent reward-based preferences

In Study 1, when predicting preferences for post-mood induction options over pre-mood induction options, there was no main effect of mood induction overall [t(33) = 0.27, *p* = 0.79] but the predicted interaction between mood induction and hypomanic traits was confirmed [Hypothesis 2b; t(33) = 2.26, *p* = .03]. Specifically, participants with elevated hypomanic traits showed a stronger mood-dependent bias on their preferences. This interactive effect replicated when predicting the continuous preference rating given for the between-block high-reward probability stimulus pair (i.e., the high-probability options learned before vs after mood induction, rated 0-100) [t(33) = 2.43, *p* = .02]. Elevated hypomanic traits were associated with a stronger mood-dependent shift in preference toward the option learned under the more positive mood state.

In Replication study, there was a trend for the mood manipulation to bias preferences for the high reward probability pair as in Study 1 (β = 5.09, *p* = 0.066), which was significant when additionally accounting for individual difference in the efficacy of the online mood induction (change in happiness, average B2-B1; β = 18.04, *p* = 0.016). However, neither a broader bias across all between-block choices nor interactions with hypomanic traits were observed (all p ≥ 0.59).

## COMPUTATIONAL MODELLING

### Simulations of the mood-biased reward integration model reproduce the empirical confidence dynamics

Following the empirical results from Study 1, we simulated a standard (unbiased) Bayesian learning model as well as two mood-biased variants in which mood was allowed to bias momentary confidence directly, or indirectly, via reward probability accumulation (Figure 3). The indirect model (Moody Accumulation; Figure 3c) yielded dynamics that matched the empirical dynamics, in which the effect of mood on confidence emerged gradually, towards the end of the learning block (Figure 3b, right). In contrast, the direct model (Moody Bonus) yielded dynamics in which confidence was maximally biased by mood early in the learning block, when the mood manipulation had maximal effect, and then tapered off (Figure 3b, left).

### Model comparison supports mood-biased accumulation of reward probability as a common mechanism for choice and confidence biases

All models in which confidence varied dynamically, as a function of trial-wise difference between stimuli pair reward probabilities, outperformed a “null confidence” model in which confidence was not dynamically updated and instead fit using linear scaling parameters. When comparing the non-moody dynamic confidence model in which mood was assumed to be static to models in which mood was allowed to vary dynamically and additionally allowed to bias confidence, the model in which mood impacted confidence via biased reward probability accumulation (Moody Accumulation) was best supported and surpassed the model in which mood directly impacted confidence (Moody Bonus) across both experiments (Figure 4). Model recovery confirmed that the Moody Accumulation model was accurately detected and distinguished from all competing models, including Moody Bonus (Supplementary Figure 4). Moreover, individual fits of the dynamic models to subjective confidence were good (median R^2^ = 0.23–0.26; representative fits shown in Figure 5b; corresponding plots for replication sample in Supplementary Figure 3) but only the Moody Accumulation model captured the divergence in confidence between positive and negative mood conditions at the end of the post-mood manipulation block (Figure 4).

**Figure 4.**
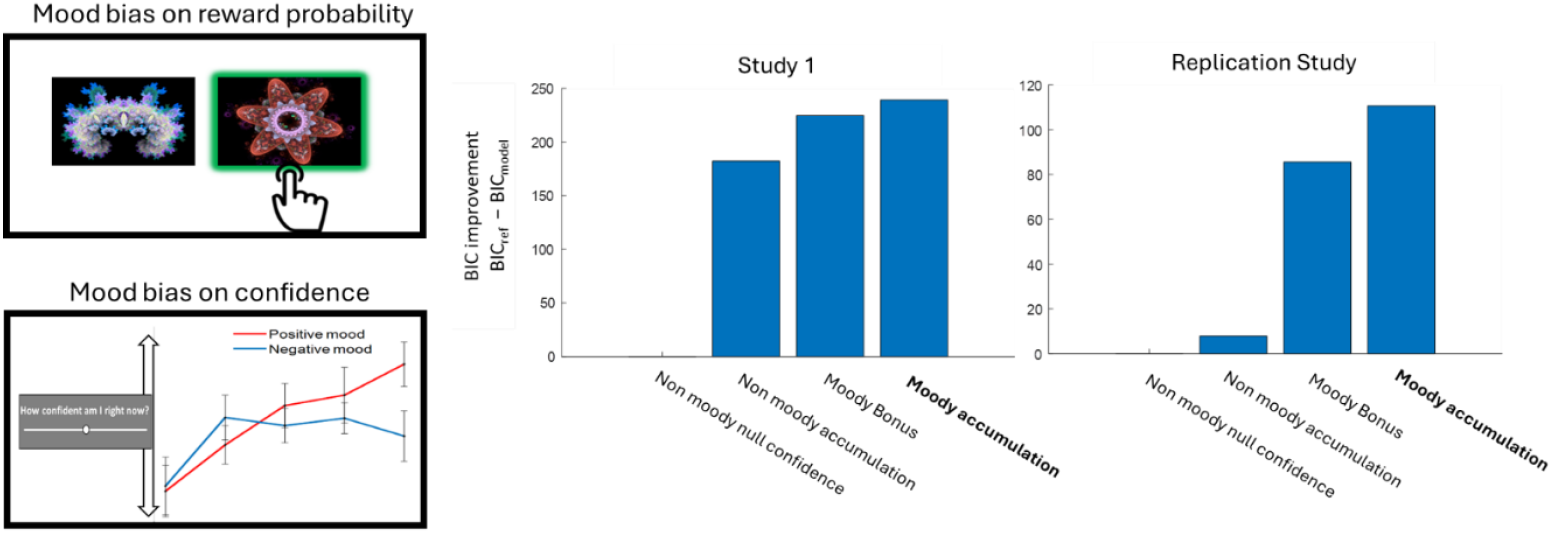
Computational model jointly capturing mood biases in reward learning and confidence. **Left panel:** The two effects are jointly captured by the model: (1) biased preference toward stimuli learned under positive relative to negative mood, and (2) mood-biased momentary confidence. **Right panel:** Model comparison results showing that, across both experiments, the recursive *Moody Accumulation* model outperformed non-moody alternatives and was the best-supported model in both studies. *Note:* The y-axis shows BIC improvement, defined as BIC_reference_ – BIC_model_; positive values indicate improved model fit relative to the reference model.

**Figure 5.**
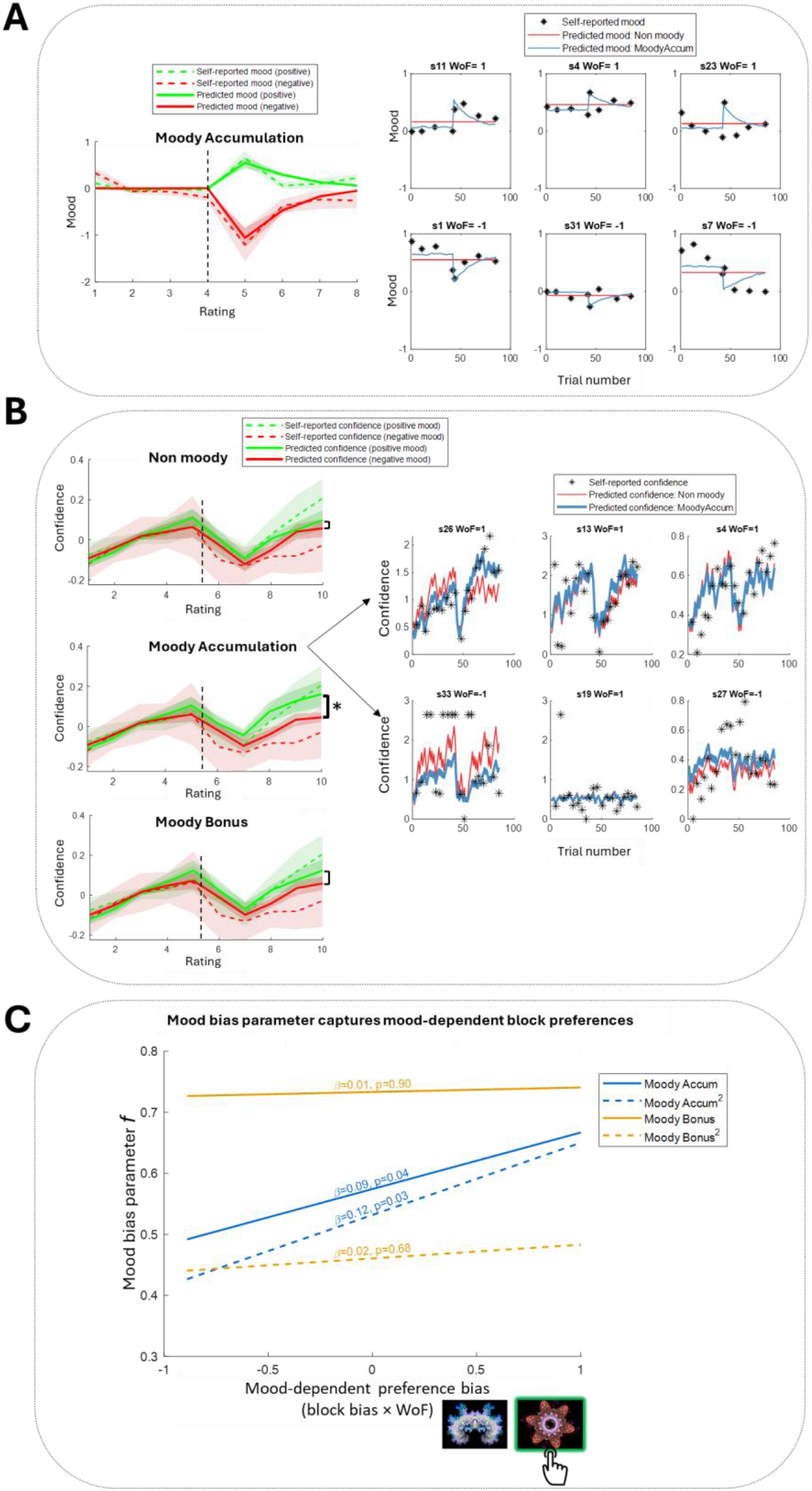
Model fits to mood, confidence, and mood-biased preferences. **A)** Model-predicted mood accorded well with self-reported happiness across all models (Moody Accumulation: median R^2^ = 0.13–0.14 for standard and quadratic model variants; Moody Bonus: median R^2^ = 0.12, both variants). Exemplar participant fits shown for good, moderate, and poorer fit (columns), separately for positive (top row; WoF = +1) and negative (bottom row; WoF = −1) mood manipulations. Dashed vertical lines separate pre- and post-mood manipulation learning blocks. **B)** All models broadly captured confidence dynamics, yet, consistent with the formal model comparison (Figure 4), only the Moody Accumulation model (median R^2^ = 0.260) captured the divergence in confidence between positive and negative mood conditions at the end of the post-mood manipulation block (asterisked timepoints), which was absent in the Non Moody (median R^2^ = 0.230) and Moody Bonus (median R^2^ = 0.240) models. Exemplar participant fits shown for good, moderate, and poorer fit (columns), separately for positive (top row; WoF = +1) and negative (bottom row; WoF = −1) mood manipulations. Group-average trajectories show binned ratings across consecutive pairs. **C)** The mood bias parameter (log *f*) from the Moody Accumulation models — but not the Moody Bonus model — predicted the manipulation-consistent preference bias (block bias × WoF; preferring positive over neutral stimulus following positive mood manipulation; neutral over negative stimulus following negative mood manipulation). Notably, these preferences are continuous slider ratings (0-100) held out from model fitting, which used only binary choices, demonstrating that the model captures a latent bias that generalises beyond the choices it was fitted to.

### A single mood bias parameter captures this shared accumulation mechanism across choice and confidence

A single mood bias parameter (*f*, Moody Accumulation model) jointly captured both mood-biased confidence (Figure 6a) and mood-biased between-block preferences at the end of the experiment (Figure 5c), supporting our hypothesis that both phenomena are underpinned by a common mechanism (Hypothesis 3a). Specifically, individuals with stronger mood bias parameters showed greater modulation of confidence post mood manipulation – higher under positive mood and lower under negative mood [*f* predicted by WoF × B2-averaged confidence interaction; Study 1: β = 0.76, t = 2.15, p = 0.04; Replication study: β = 0.48, t = 2.31, p = 0.02; Figure 6a]. This result was robust to operationalisation of confidence, holding when using change in confidence (B2 – B1, i.e. the averaged increase or decrease following mood induction; *f* predicted by ΔConfidence × WoF; p ≤ 0.014).

**Figure 6.**
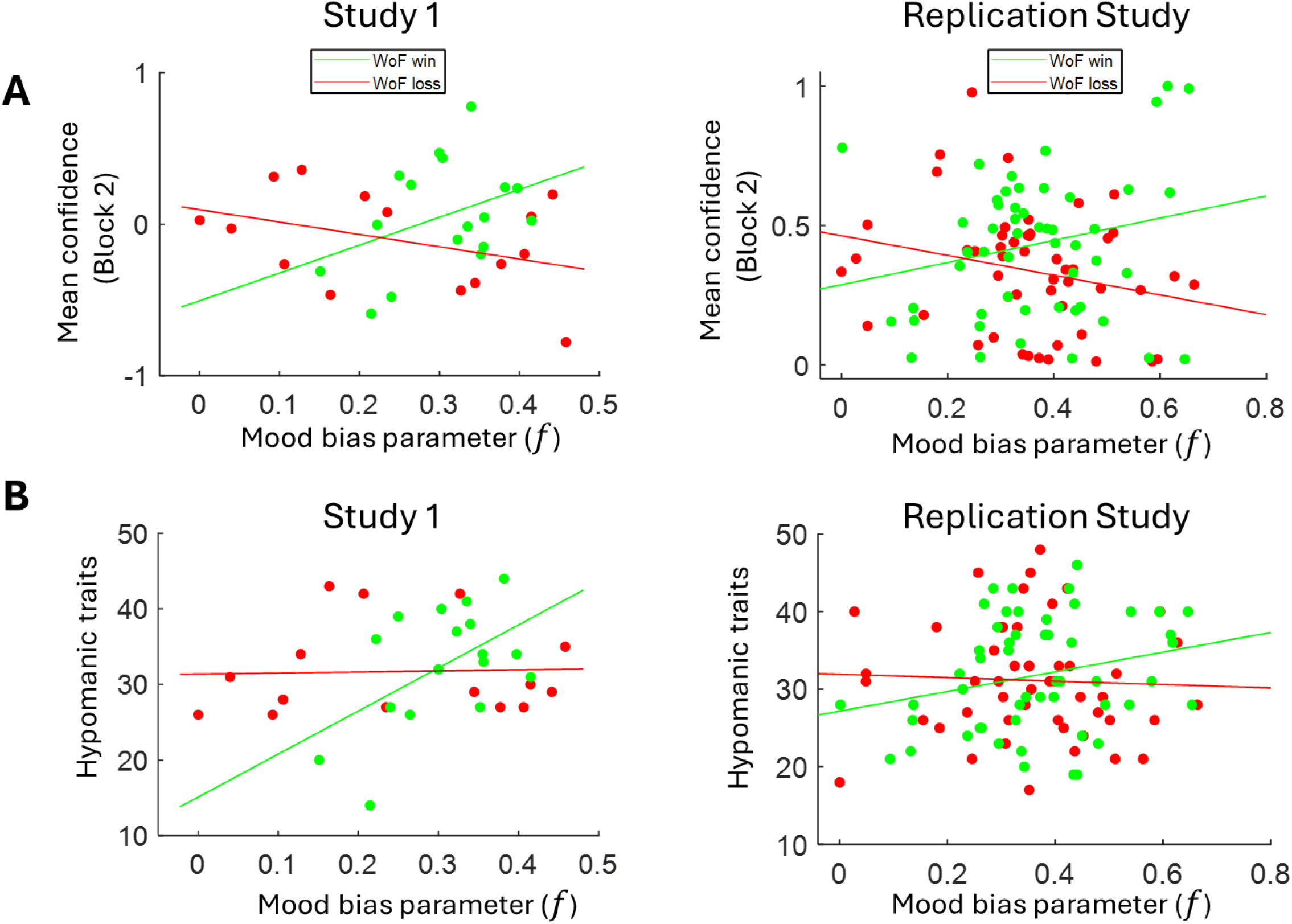
Relationship between mood bias parameter, the effect of mood on confidence and individual differences in hypomanic traits. **A)** In both experiments, the mood bias parameter (log *f*) from the winning Moody Accumulation model was correlated with the extent that mood biased confidence during the post-mood manipulation learning block. Specifically, higher *f* parameter accounted for the empirically observed overconfidence under positive mood (WoF win; green markers), and under-confidence under negative mood (WoF lose; red markers) [WoF × confidence interaction; Study 1: β = 0.76, t = 2.15, p = 0.04; Replication study: β = 0.48, t = 2.31, p = 0.02]. **B)** Under positive mood, higher hypomanic traits were predicted by a stronger mood bias parameter [Study 1: r = 0.53, p = 0.029; Replication study: r = 0.32, p = 0.02].

As a further robustness check, given prior work indicating that confidence is best captured as the squared difference in value and probability ^19,28,29^, we compared model variants in which confidence was formalized as the squared difference between stimulus Q values. The quadratic Moody Accumulation (Moody Accumulation quadratic) variant did not outperform the non-quadratic model in Study 1 (but was marginally better supported in the Replication study; Supplementary Figure 5) and still captured the effect of mood on confidence in the mood bias parameter (p = 0.019).

### Mood-biased accumulation of reward probability is amplified in individuals with higher hypomanic traits

Finally, we confirmed that the mood bias mechanism also underpinned the non-modelling findings that higher hypomanic traits are associated with stronger effects of mood on confidence. Across both experiments, stronger mood bias parameter (*f*) predicted higher hypomanic traits under positive mood (Figure 6b). In Study 1, *f* interacted with mood manipulation [β = 0.46, t = 2.13, p = 0.04], with stronger mood bias under positive than negative mood [positive: r(16) = 0.53, p = 0.029; negative: r(15) = 0.035, p = 0.9]. Both results replicated in the Replication study [r(53) = 0.32, p = 0.02; negative: r(49) = −0.046, p = 0.76].

## Discussion

Our goal was to identify the computational mechanism through which mood influences confidence during learning, and to test whether this process was amplified in individuals prone to instability in mood and self-efficacy. This question is particularly relevant for mood disorders such as bipolar disorder, which have been linked to dysregulated reward sensitivity and disrupted self-appraisal^18^. In recent work, we have shown that individuals with bipolar disorder exhibit exaggerated mood-momentum biases in striatal reward prediction error signals^7^, consistent with a neural implementation of recursive mood-learning mechanisms. Here, we extend this framework to investigate how mood shapes confidence. Across two independent datasets—one laboratory-based and one pre-registered online replication—we show that mood biases confidence not through a direct shift in self-belief, but by distorting how individuals learn about reward contingencies. We formalised this account in a computational model in which mood recursively biases the accumulation of reward probability estimates, which in turn shape subjective confidence. This model jointly captured the impact of mood on learning and confidence, and explained the amplification of these effects in individuals with elevated hypomanic traits, which confer instability in mood and self-efficacy^9,14^.

Across both studies, the mood manipulation reliably altered subjective happiness, with positive and negative mood manipulations increasing and decreasing mood, respectively (Figure 3b, left). These shifts in mood biased participants’ learning in two key ways. First, participants expressed greater preference for stimuli encountered under positive mood, indicating that mood influenced the perceived value of rewards. Second, confidence ratings diverged following mood induction—participants in a positive mood became increasingly confident across trials, while those in a negative mood showed the opposite pattern—despite equivalent objective accuracy (Figure 2, left). Importantly, this effect of mood on confidence accumulated gradually with learning, rather than emerging immediately after the manipulation when mood change was maximal (Figure 3b, right), suggesting that the effect emerges in concert with reward integration processes during learning. This temporal dissociation between mood and confidence provides empirical support for a recursive model in which mood exerts its influence not directly, but by biasing the encoding of reward outcomes and the updating of internal value estimates. Computational modelling supported this interpretation. Among the candidate models, the recursive Moody Accumulation model—where mood biases confidence indirectly by altering the integration of reward outcomes—outperformed alternatives across both datasets. This model captured the gradual emergence of mood-biased confidence over time (Figure 3c), aligning with the empirical dynamics observed in participants (Figure 3b, right). In contrast, a model in which mood directly shifted confidence (Moody Bonus) failed to reproduce these dynamics, predicting instead a transient effect peaking immediately after mood induction. Crucially, the *Moody Accumulation* model’s mood bias parameter, jointly accounted for both key behavioural effects: the mood-driven shift in confidence and the preference for stimuli learned under positive mood (Figures 5c, 6a).

Consistent with theories of mood as representing environmental momentum^2^, these findings suggest that mood dynamically distorts reward prediction error signals and resulting value representations that underpin both decision-making and momentary confidence. In keeping with the theory, this mechanism appears normative, operating across participants regardless of trait vulnerability. Previous neuroimaging studies in non-clinical samples have identified confidence-related signals in ventromedial prefrontal cortex and striatum during reward-based decisions^19,29^, regions that also encode reward prediction errors biased by mood^5–7^. This overlap supports the idea that mood-biased confidence reflects altered input to these valuation systems, with mood-biased RPEs propagating downstream to shape confidence estimates.

In both studies, individuals with elevated hypomanic traits showed greater susceptibility to mood-biased confidence — overconfident under positive mood states and underconfident under negative mood states, relative to objective accuracy (Figure 2). This represents an amplification of the normative (group-level) effect observed in both studies. This stronger mood bias was recapitulated at the computational level, which predicted hypomania traits specifically under positive mood (Figure 6b). This asymmetry suggests that elevated hypomanic traits more strongly amplify mood-biased learning under positive mood, which would be in keeping with findings of pervasive reward hypersensitivity^10–13^ and the focus of the Hypomanic Personality Scale on propensity for elevated mood.

Confidence in this task can be interpreted in at least two ways. At one level, it reflects a metacognitive evaluation of decision accuracy—how certain participants are that their choice will yield reward. At another, it may reflect a proximal sense of control or action utility, related to the experience of agency. These perspectives are not mutually exclusive, and both may be relevant to mood disorder symptoms. Notably, mood in our study biased confidence without affecting objective accuracy, echoing findings from Rouault and colleagues^22^, who showed that psychiatric symptoms were associated with distorted metacognitive insight despite preserved accuracy. These results raise the possibility that apparent shifts in confidence across mood states—heightened during (hypo)mania or diminished in depression—may not reflect irrational beliefs per se, but rather rational inferences drawn from systematically biased learning signals. This suggests that mood biases the evidence upon which confidence is formed, leading to judgments that appear irrational but are internally consistent given the distorted inputs. Future work could test whether mood fluctuations preferentially bias specific levels of the confidence hierarchy, and whether distortions at different levels contribute uniquely to symptoms such as poor insight, grandiosity, and unstable self-esteem across mood states.

Our design offers several strengths. Most notably, it tracked the evolution of mood-biased confidence in real time, rather than relying on retrospective judgments or trait-level self-report, capturing dynamic fluctuations directly linked to trial-by-trial learning processes. Crucially, these confidence shifts were independent of objective accuracy, consistent with mood biasing the interpretation of outcomes rather than performance itself. This aligns with clinical observations that mood episodes are accompanied by distorted evaluations of reward and action utility^30,31^ and builds on prior evidence linking reward sensitivity and disrupted goal pursuit to affective instability^32,33^.

Nonetheless, several limitations should be noted. First, while individual differences in hypomanic traits consistently amplified mood biases in confidence, replicating across both datasets, we partially replicated the prior finding that hypomanic traits strengthen mood biases on the reward-based preferences sampled at the end of the experiment^5^. We observed this result in our laboratory study, whereas in our online replication, mood effects on preference were present but not further moderated by hypomanic traits. Notably, however, computational modelling results converged across both datasets and this prior study, that found hypomanic traits were associated with a stronger mood bias parameter^5^. End-of-experiment preference measures provide a brief and distal readout of accumulated learning and may depend more strongly on task engagement, which is inherently more variable in online settings^34^. By contrast, the computational model additionally leveraged trial-by-trial choices and confidence ratings throughout learning, which may explain its greater sensitivity to individual differences in hypomanic traits across datasets. Confidence ratings, being more proximal to ongoing learning, may provide a particularly sensitive readout of mood biases in future studies.

A second limitation concerns the operationalisation of confidence in our task. Recent theoretical frameworks distinguish between multiple levels of confidence, ranging from trial-by-trial decision certainty to broader self-beliefs about skills and abilities^26^. Accordingly, our task indexed momentary confidence in which option was more likely to yield reward—that is, confidence in the relative reward probabilities of the available actions—and we deliberately sampled this after outcome feedback, when mood could impact value updating and thus exert its strongest influence on perceived efficacy. This approach was motivated by prior work linking hypomanic traits to elevated success expectancy following reward ^14^. In contrast, other paradigms typically assess confidence at the time of choice (e.g.^19,22,29^), which may yield estimates less influenced by recent outcomes and more decoupled from momentary mood. While our approach allowed us to capture dynamic interactions between mood, learning, and confidence, it also means that our confidence measure may reflect a blend of metacognitive certainty and transient affective states. However, we ruled out the possibility that confidence simply tracked mood. First, confidence and mood showed dissociable temporal dynamics (Figure 3b, right), including in a separate block with dense mood sampling (Supplementary Figure 2). Second, the mood effect on confidence persisted when controlling for recent outcomes (Supplementary Table 2), suggesting that mood-biased confidence cannot be fully explained by feedback alone. We also tested alternative model specifications for the mapping between subjective reward probability and confidence. While our linear model fit both datasets well, we also tested a quadratic variant used in other studies of confidence ^19,28,29^ which performed marginally better in the online sample. Crucially, the core mood bias effect was robust across both model variants.

Finally, although the Hypomanic Personality Scale captures trait vulnerability to bipolar disorder^8,9^, future work in clinical populations is needed to confirm whether the mood-biased confidence effects reported here extend to individuals with bipolar disorder, where analogous biases in striatal reward prediction error signals have already been established^7^. Neural validation would be valuable, particularly in striatal and prefrontal regions implicated in both mood regulation and confidence estimation, to test whether mood-biased RPEs propagate downstream to distort confidence as proposed.

In conclusion, we provide evidence for a unified model in which mood shapes confidence dynamically, through biased accumulation of reward probability estimates during learning. This mechanism offers a principled account of how transient emotional states can give rise to persistent distortions in decision-making and confidence, particularly in individuals prone to instability in mood and self-efficacy. By linking mood, learning, and confidence within a formal computational framework, our findings offer a new lens through which to understand the cognitive distortions seen in mood disorders — not as direct consequences of mood or belief change, but as emergent properties of mood-biased reward learning.

## Methods

### Participants and Measures

In Study 1, 35 participants were recruited from the UCL student population via online advert and participated in person, at the laboratory. The Replication study was powered (G*Power) to achieve 95% power to detect the effect sizes from Study 1: the effect of mood on biased confidence (Hypothesis 1a; d = 0.71) and its interaction with hypomanic traits (Hypothesis 1b; d = 0.82). Consequently 106 participants were recruited through Prolific (https://www.prolific.com). Study design, predictions and analyses were pre-registered (https://osf.io/ygc4t; Supplementary Table 1). Participants completed the International Personality Item Pool^35^ version of the Hypomanic Personality Scale which captures marked shifts in mood^36^ and confidence^14^ and is an established psychometric risk factor for bipolar disorder^8,9^.

### Task

#### Study 1

We used a previously validated task^5^ to quantify the influence of mood on reinforcement learning (Figure 1). The task consisted of two blocks, in which participants chose between a pair of stimuli with different probabilities of reward (33% and 66% of receiving £0.10), with one block before and one after a mood manipulation. There were 42 trials per block, with stimuli alternating their presentation side (left/right) pseudo-randomly. Ten of these trials were forced choice, in which only one of the options was available to choose, to facilitate comparable estimates of the reward probabilities of all stimuli. A new stimuli pair was learned in the second block but, crucially, these had the same ground truth probability of reward as in block one (i.e., 33% and 66%). The mood manipulation consisted of spinning a wheel of fortune in which participants received either a large monetary gain or loss of £5, yielding two groups (positive mood group and negative mood group). Both mood and confidence were measured periodically throughout the learning blocks by self-report. Participants gave periodic ratings of their confidence and were instructed to rate this as the degree of certainty at any given time about which of the available options was the option that yields more frequent reward. During the experiment they were prompted (“How confident do you feel right now?”, every 4-5 trials, totalling 10 ratings per block). Mood was sampled three times during the learning block, in the first, second and final third of trials (“How happy do you feel right now?”). To confirm that confidence and happiness are separable phenomena that follow distinct dynamics during learning, we collected a supplementary learning block with new stimuli (Supplementary Figure 2), in which participants were instead predominantly asked to rate mood (10 ratings per block) and sporadically their confidence (3 ratings per block).

At the end of the task, participants were asked to indicate preference in pairwise choices between all the stimuli learned throughout the experiment. Each pairwise choice was repeated once, with participants this time giving a slider rating of preference (converted to −50 to +50, with 0 being indifferent; no preference between the two stimuli). The within-block pairs give a readout of recall accuracy (which was the higher reward option from each pair faced). The between-block pairs give a read-out of the extent that mood biased reward learning during the task^5^.

#### Replication study

The above task was administered online via the *Prolific* platform and followed the same design, with an increase in trial number from 42 to 46 trials per block including 8 forced choice trials to offset potential differences in attention and task engagement in online settings.

### Planned analyses

Pre-registered predictions (Supplementary Table 1) were tested using general linear models, linear mixed-effects analyses, and computational modelling. We predicted that mood would bias subjective confidence independently of objective accuracy (Hypothesis 1a), and that this effect would be stronger in individuals with higher hypomanic traits (Hypothesis 1b). We additionally predicted that mood would bias reward learning, indexed by post-induction stimulus preferences (Hypothesis 2a), with stronger effects in those with higher hypomanic traits (Hypothesis 2b). Finally, we predicted that a recursive model in which mood biases the accumulation of reward probability estimates — rather than directly shifting reward expectancy — would best account for the data (Hypothesis 3a), with a single mood-bias parameter jointly capturing effects on both confidence and preferences (Hypothesis 3b).

To test for the trial-by-trial impact of mood on confidence, trial-level data were entered into linear mixed effects models. Confidence reports were predicted with fixed effects of the mood manipulation received, as well as objective accuracy (the cumulative proportion of choices of the high reward option) and hypomanic traits. Random effects were participant ID, with separate intercept and slope for trial-wise accuracy:

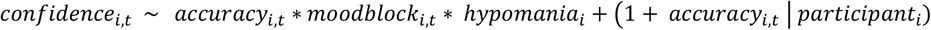

To test whether mood biased reward learning (Hypothesis 2), we examined pairwise binary preferences for stimuli encountered post-versus pre-mood induction using regression analyses, testing both the main effect of mood manipulation received and its interaction with HPS score^5^. We corroborated this with analysis of the continuous preference rating for the between-block high-reward probability pair, given that these stimuli were encountered more frequently during learning, providing better-sampled Q value estimates and greater opportunity for mood-biased evidence accumulation to manifest.

The remaining pre-registered predictions (Supplementary Table 1) were tested using computational modelling: that a recursive learning model in which mood biases the accumulation of reward probability estimates would be best supported by the data (Hypothesis 3a); and that a single mood-bias parameter within this model would account for mood-related effects on both subjective confidence and reward learning (Hypothesis 3b).

After removing participants with incomplete data (Study 1: 1 participant, Replication study: 4 participants), the specified, directional hypotheses above were tested at conventional significance threshold of 0.05.

### Computational model of mood and confidence

Building on the approach of Eldar & Niv^5^ and given our focus on confidence, we specified Bayesian learning models in which reward probabilities are computed based on available reward evidence, with confidence formalised as the trial-wise difference in Q values between available options. The details of the models tested are in Table 2. The test choice trials were included in the modelling in Study 1, but omitted in the replication study, as mood effects on preferences were present only in the continuous preference ratings and not binary choices fitted by the model.

**Table 2.**
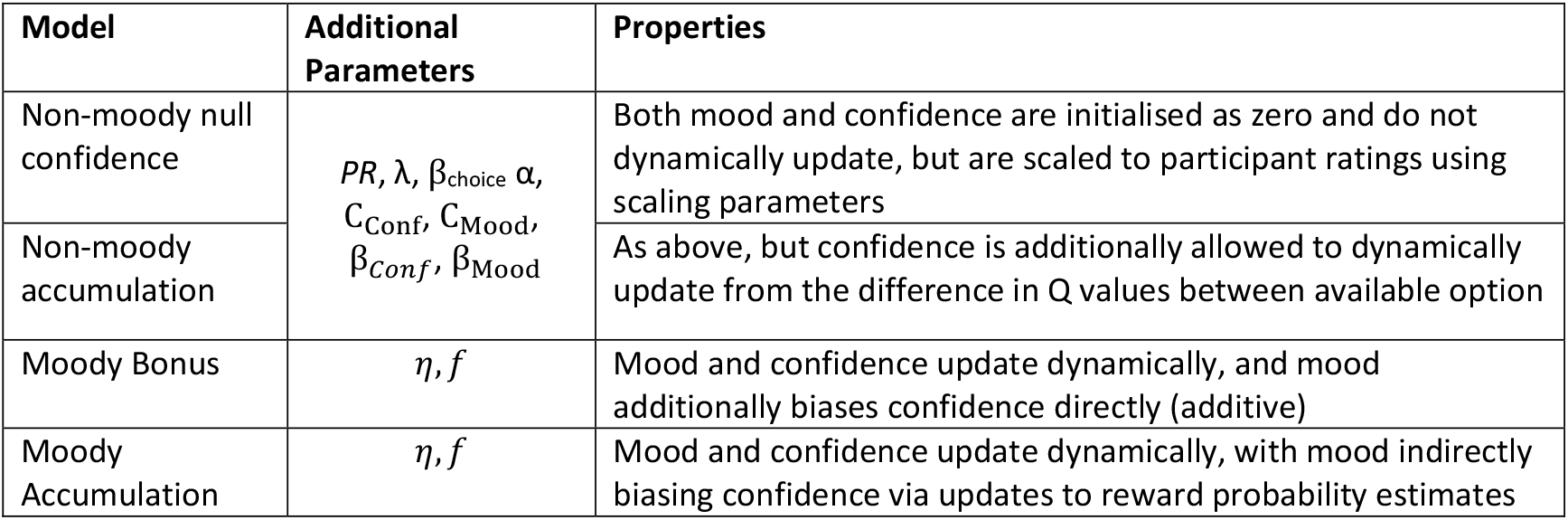
Description of computational models. The base non-moody models consist of the following parameters: strength of prior about reward probability (*PR*), forgetting back to prior (λ), a choice inverse temperature (β_choice_), a Q value mixing parameter (α), plus parameters that centre and scale mood (C_Mood_, β_Mood_) and confidence (C_Conf_, β_Conf_) to participant ratings. In addition to these core parameters, the moody models both included a mood update rate (*η*) and, in the Moody Accumulation model, a mood bias parameter (*f*).

#### 1. Learning of reward probabilities

Reward probability was allowed to accumulate under a beta-distribution learning rule:

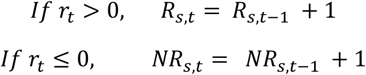

Under the recursive Moody Accumulation model, mood (*m*) was allowed to bias the impact of reward observations on learning through a mood bias parameter (*f*) and resulting mood-dependent gain term, *g*_*t*_. This gain term modulates the influence of reward outcomes on value updating in a mood-dependent manner, with the direction of the effect determined by the sign of *f*:

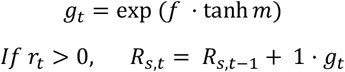

The mood bias parameter *f*was drawn from a zero-mean Gaussian prior, allowing for bidirectional coupling between mood and reward integration. Positive values of *f*produce canonical mood-dependent effects, whereby positive mood amplifies and negative mood attenuates the impact of reward outcomes on value updating. In contrast, negative values of *f*permit an inverted or paradoxical coupling, in which positive mood reduces, and negative mood enhances, the influence of reward outcomes on subsequent value estimates. All models included a uniform prior about reward probability, whose strength (*PR*) served to slow down learning from small numbers of observations. A forgetting parameter, λ, drawn from a beta distribution, allowed the reward probability of the chosen option to decay back to the prior:

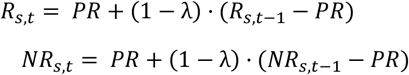

On each trial, Q values were computed. We reasoned that participants might track two quantities relevant to reward probability to varying degrees (evidence strength: an estimate of how many more rewards the current option has given than non-rewards, as well as the estimated probability of reward):

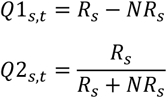

We allowed the degree to which each participant was tracking each quantity to vary, through a mixing parameter, *α*:

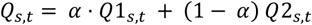

The probability of choosing stimulus *i* at trial *t* was controlled by an inverse temperature parameter (β_choice_):

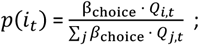

where *j* denotes the stimuli available at trial *t*.

#### 2. Formalising of mood and confidence

Mood on each trial was computed by first taking the Q value update for the chosen option as the reward prediction error term. Although reward prediction errors and value updates sometimes diverge, evidence^37^ and theory^38^ indicate that the latter quantity offers a better explanation for mood. Supplementary model comparison confirmed that a unified mixing parameter for the Q values used for choice, the Q values used for confidence and reward prediction error (RPE) updates was better supported than separate mixing parameters for each quantity.

Then, mood is updated based on the RPE (*δ*), scaled by a mood update rate parameter, *η*:

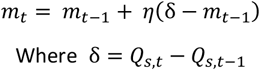

Model estimates of confidence were operationalised as the difference between the Q values of the two available options on each trial. We then fit trial-wise model estimates of confidence (*c*_*t*_) and mood (*m*_t_) to their respective subjective ratings given by participants (after inverse hyperbolic tangent transformation to account for the bounded nature of ratings, using constant (C_Conf_, C_Mood_) and scaling (β_C_, β_M_) parameters:

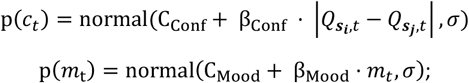

where *σ* is a free parameter denoting the standard deviation of reported values from model predictions.

Under the Moody Bonus model, mood does not alter learning of reward contingencies, but instead rescales the separation between already learned value beliefs, amplifying this separation under positive mood and compressing it under negative mood. Consequently, the same form of mood-dependent gain term as in the Moody Accumulation model, *g*_*t*_, was applied directly to the trial-wise confidence quantity, before including in the above likelihood function. When this term is augmentative (i.e., greater than 1), it increases the magnitude of the Q value difference on that trial; when it is diminutive (i.e., less than 1), it compresses the magnitude of the Q value difference, before scaling by *β*_*C*_ and *β*_*M*_:

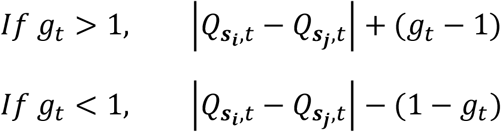

### Model fitting procedures

Model fitting followed the approach of Eldar & Niv (2015)^5^, using importance sampling to approximate model evidence as the mean log likelihood across 100,000 random parameter samples. Parameters were sampled from weakly informative priors chosen to reflect parameter constraints: bounded parameters (λ, *η*, α) from Beta(1,1) uniform priors; positive-only parameters (*PR*, β_choice_) from Gamma(1,1); and unbounded parameters for mood and confidence scaling and *f* from Normal(0,1). Model comparison used the Bayesian Information Criterion (BIC), approximated as −2 × log mean likelihood + k × log(n), where k is the number of free parameters and n is the number of observations, with lower BIC indicating better fit. BIC improvement over a reference model (BICreference − BICmodel) is reported to facilitate comparison across models with different numbers of parameters.

### AI tools statement

AI-assisted writing tools were used in the preparation of this manuscript. All content was reviewed and approved by the authors.

## Acknowledgements

This work was supported by a Medical Research Council Clinician Scientist Fellowship awarded to L.M. (MR/S006613/1).

## Author contributions

L.M. conceived and designed the study, analysed the data, and wrote the manuscript. S.P.W. collected the data and contributed to data analysis. E.E. and R.B.R. advised on analysis and contributed to manuscript preparation. All authors reviewed and approved the final manuscript.

## SUPPLEMENTARY MATERIAL

**Supplementary Table 1.** Pre-registered hypotheses and mapping to manuscript hypotheses and corresponding results section.

| Preregistered predictions | Prediction | Manuscript hypothesis label | Corresponding results section |
| --- | --- | --- | --- |
| 1 | The mood induction would be effective in impacting mood, with winning on the wheel of fortune increasing happiness and losing on the wheel of fortune decreasing happiness. | [Manipulation check] | Results: *Confirmation of learning and mood manipulation* |
| 2 | The resulting change in mood would bias subjective confidence, independent of objective performance, such that increases in happiness would increase confidence and decreases in happiness would decrease confidence. | Hypothesis 1a | Results: *Mood biases momentary confidence, amplified by trait instability in mood and self-efficacy* |
| 4 | Effects of mood on confidence [and on reward learning] would be stronger in participants with higher levels of trait hypomania. | Hypothesis 1b |  |
| 3 | Change in mood would bias reward learning, with participants preferring stimuli encountered in the block in which they were happier. | Hypothesis 2a | Results: *Mood during learning biases subsequent reward-based preferences* |
| 4 | Effects of mood [on confidence and] on reward learning would be stronger in participants with higher levels of trait hypomania. | Hypothesis 2b |  |
| 5 | A recursive moody reward probability accumulation model [Moody Accumulation] would be best supported. | Hypothesis 3a | Results: *Model comparison supports mood-biased accumulation of reward probability as a common mechanism for choice and confidence biases* |
| 6 | Effects of mood on confidence and on learning of reward probability would be explained by a single mood bias parameter. | Hypothesis 3b | Results: *A single mood-bias parameter captures this shared accumulation mechanism across choice and confidence* |

### Confirmation of learning and mood manipulation

Participants chose the high probability option more than chance in Study 1 (Block 1: mean ± SEM = 67.9% ± 0.01; t(34) = 12.1, *p* < 10^−14^; Block 2: mean ± SEM = 69.5% ± 0.02; t(34) = 10.7, *p* < 10^−13^). Participants also chose the high probability option more than chance in the replication study (Block 1: mean ± SEM = 59.2% ± 0.02; t(104) = 4.18, *p* < 10^−5^; Block 2: mean ± SEM = 57.4% ± 0.02; t(104) = 4.59, *p* < 10^−6^). The mood manipulation impacted momentary happiness in Study 1 with greater increase in happiness after the positive compared to the negative mood induction [between-groups difference in change in happiness, block2 - block 1, for positive mood minus negative mood group: t(33) = 4.21, p = 0.00019]. Follow up tests confirmed that positive mood manipulation increased happiness (mean change in happiness block 2 – block 1 ± SEM = 9.22% ± 0.02; t(17) = 4.21, p = 0.00029) whereas negative mood manipulation decreased happiness (mean change ± SEM = −14.5% ± 0.05; t(16) = 2.72, p = 0.0075). Similarly in Replication study, change in block-average happiness was higher in the positive mood induction compared to negative mood induction group [t(103) = 7.65, *p* < 10^−6^], with increased happiness after positive mood manipulation (mean ± SEM = 7.25% ± 0.02; t(54) = 3.22, *p* = 0.0011) and decreased happiness after negative mood manipulation (mean ± SEM = −20.1% ± 0.03; t(49) = 8.22, *p* < 10^−11^).

**Supplementary Table 2.**
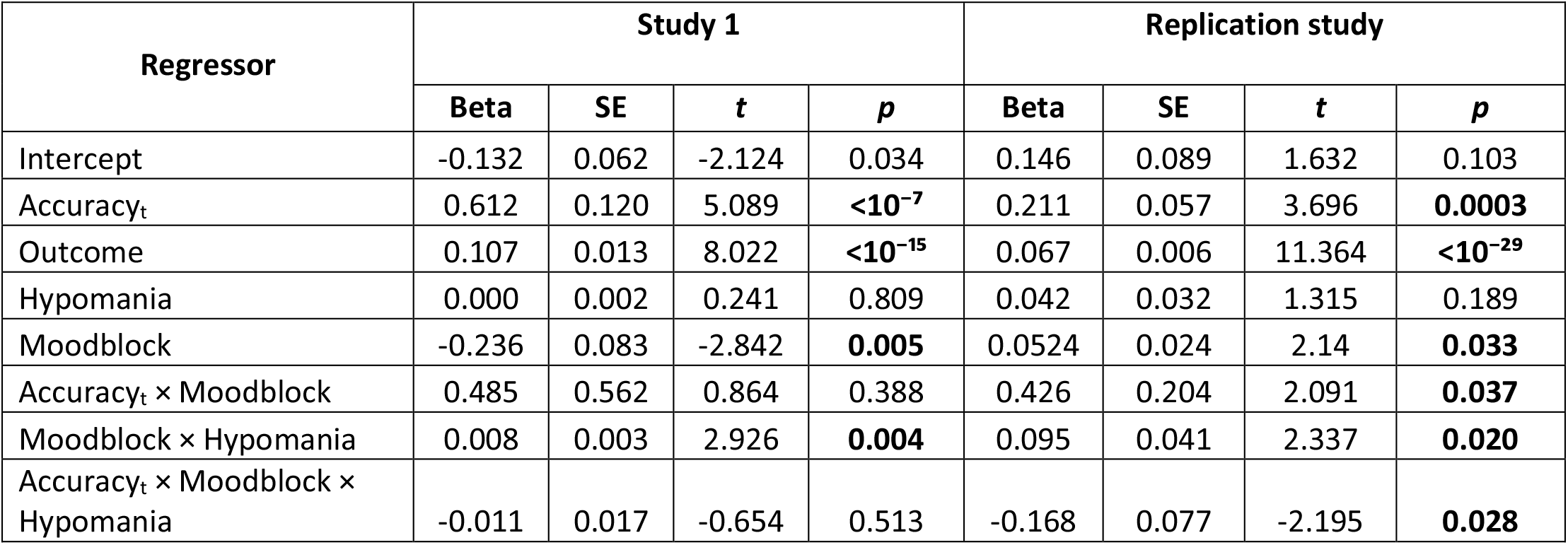
Linear mixed effects model of the influence of mood block in predicting subjective confidence over and above trial wise accuracy, additionally accounting for most recent outcome observed.

| Regressor | Study 1 |  |  |  | Replication study |  |  |  |
| --- | --- | --- | --- | --- | --- | --- | --- | --- |
|  | Beta | SE | *t* | *p* | Beta | SE | *t* | *p* |
| Intercept | -0.132 | 0.062 | -2.124 | 0.034 | 0.146 | 0.089 | 1.632 | 0.103 |
| Accuracy<sub>t</sub> | 0.612 | 0.120 | 5.089 | **<10<sup>-7</sup>** | 0.211 | 0.057 | 3.696 | **0.0003** |
| Outcome | 0.107 | 0.013 | 8.022 | **<10<sup>-15</sup>** | 0.067 | 0.006 | 11.364 | **<10<sup>-29</sup>** |
| Hypomania | 0.000 | 0.002 | 0.241 | 0.809 | 0.042 | 0.032 | 1.315 | 0.189 |
| Moodblock | -0.236 | 0.083 | -2.842 | **0.005** | 0.0524 | 0.024 | 2.14 | **0.033** |
| Accuracy<sub>t</sub> × Moodblock | 0.485 | 0.562 | 0.864 | 0.388 | 0.426 | 0.204 | 2.091 | **0.037** |
| Moodblock × Hypomania | 0.008 | 0.003 | 2.926 | **0.004** | 0.095 | 0.041 | 2.337 | **0.020** |
| Accuracy<sub>t</sub> × Moodblock × Hypomania | -0.011 | 0.017 | -0.654 | 0.513 | -0.168 | 0.077 | -2.195 | **0.028** |

**Supplementary Figure 1.**
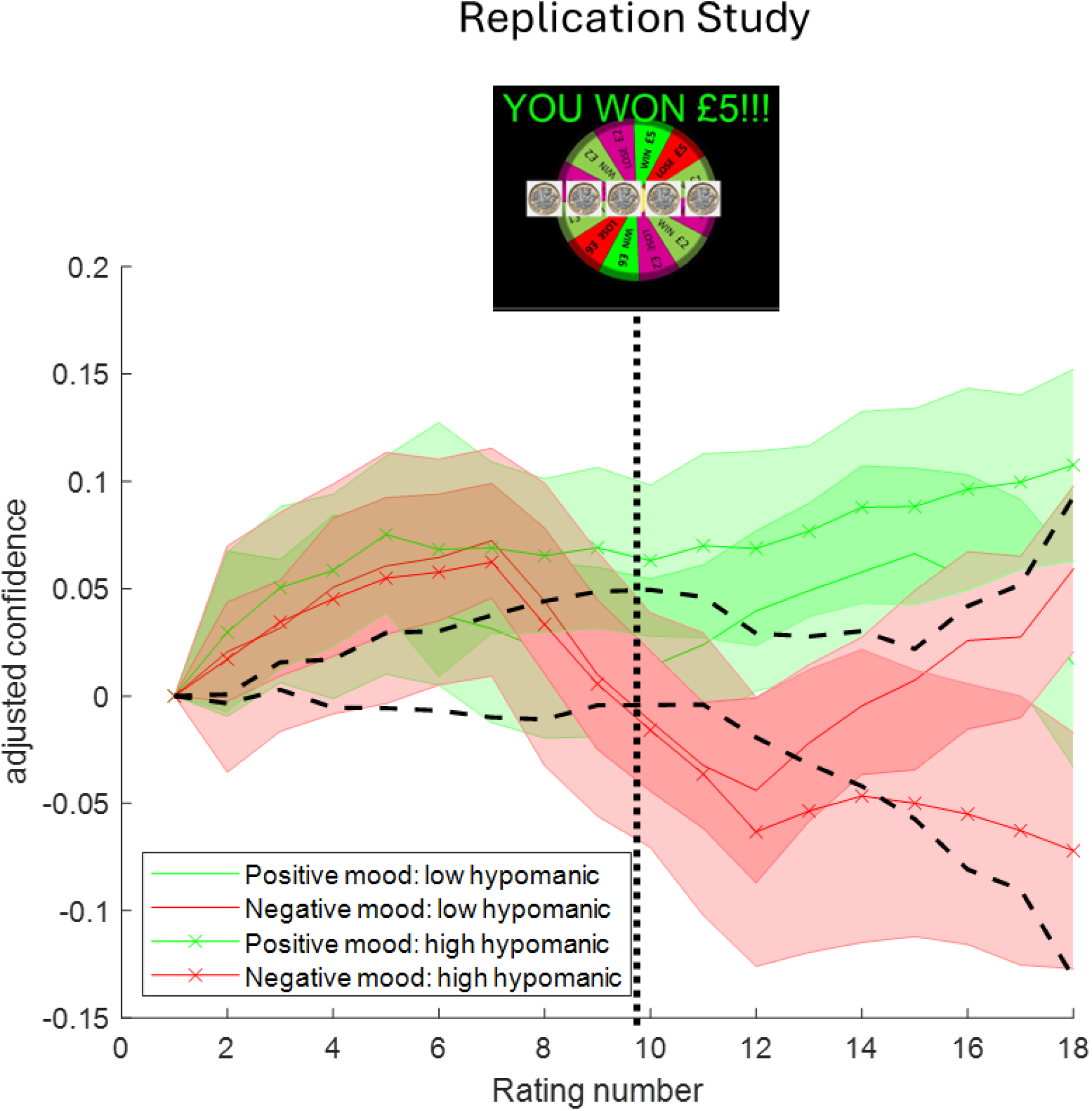
Mood biases momentary confidence and more so in those with elevated hypomanic traits. Following mood induction (vertical dashed line), confidence ratings diverge depending on mood valence, with stronger effects in participants with higher hypomanic traits (median split shown for visualisation). The difference between positive and negative mood conditions increases over subsequent learning trials, consistent with mood-dependent biases accumulating with learning. Dashed lines illustrate the stronger divergence expected in participants with higher hypomanic traits, corresponding to relative over-confidence under positive mood and under-confidence under negative mood.

**Supplementary Figure 2.**
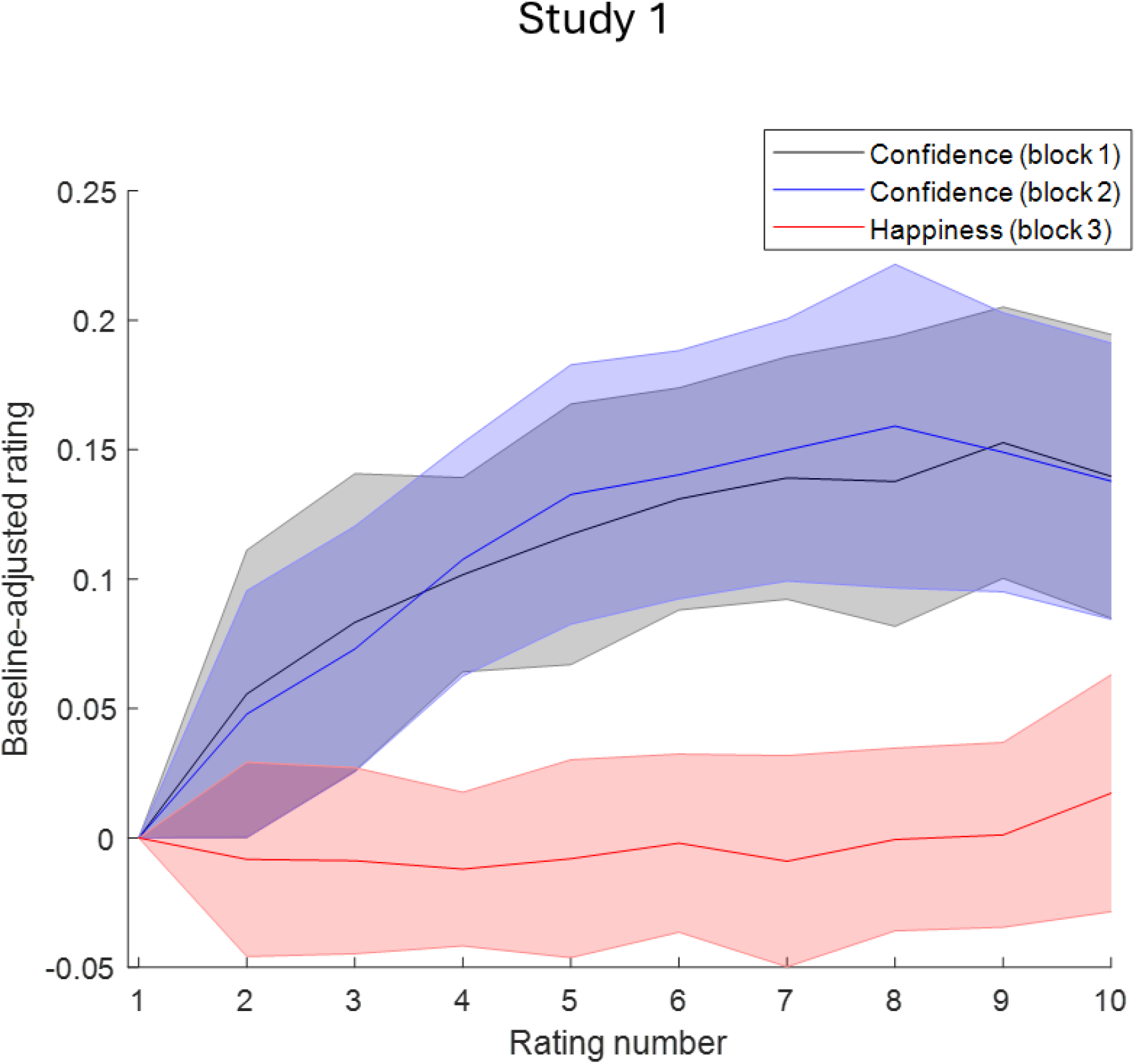
Confidence and happiness have distinct dynamics. Mean self-reported confidence (blocks 1 and 2) and happiness (block 3) in Study 1 (baseline-adjusted to each participant’s first rating in the block). Confidence increased across ratings during the learning blocks and was strongly correlated across blocks 1 and 2 (*r* = 0.74, *p* = 0.02). In contrast, happiness did not increase during the control block and was unrelated to confidence (block 1: *r* = 0.10, *p* = 0.78; block 2: *r* = 0.02, *p* = 0.96).

**Supplementary Figure 3.**
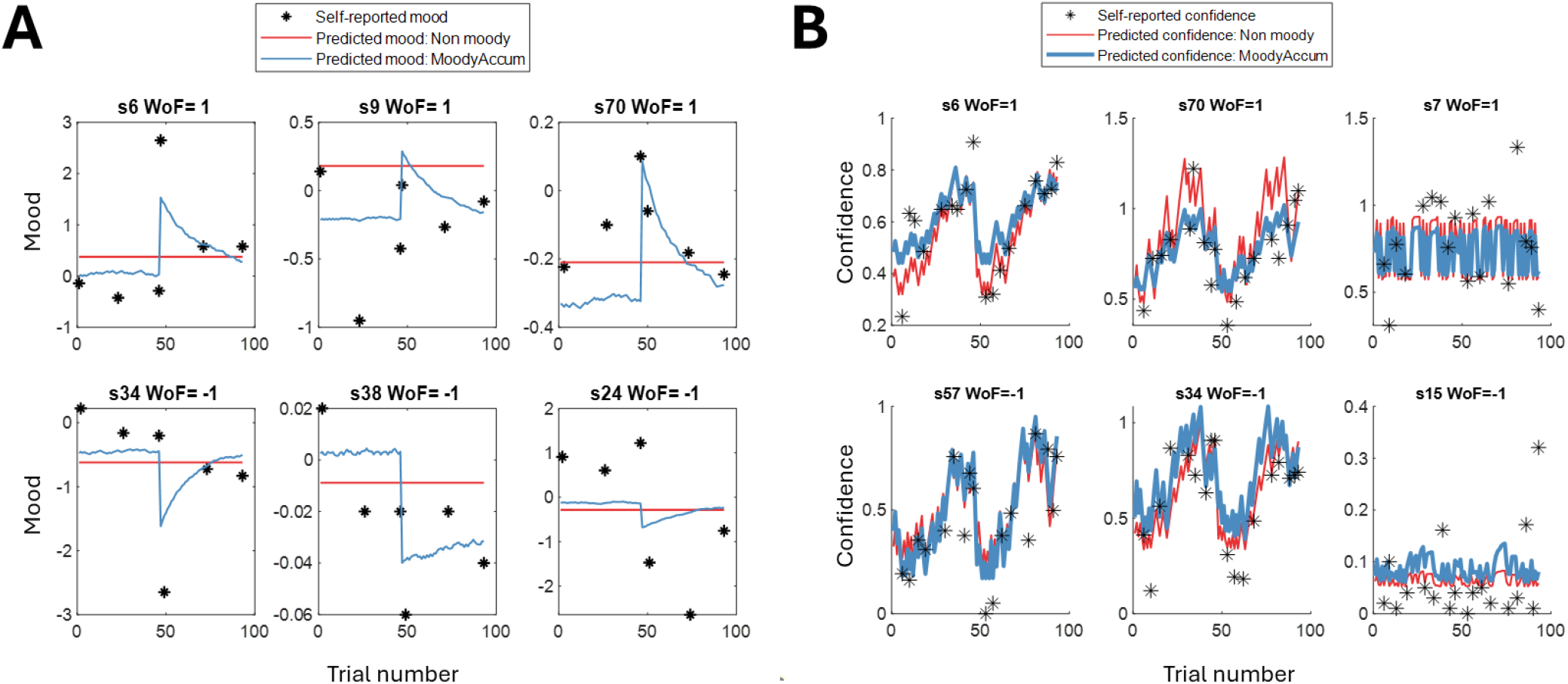
Model fits for mood and confidence in online Replication study. **A)** Model-predicted mood accorded well with self-reported happiness across all models (Moody Accumulation: median R^2^ = 0.17 and 0.19 for standard and quadratic model variants; Moody Bonus: median R^2^ = 0.2 and 0.16, for standard and quadratic model variants variants). **B)** All models broadly captured confidence dynamics (median R^2^ = 0.06–0.092). Exemplar participant fits shown for good, moderate, and poorer fit (columns), separately for positive (top row; WoF = +1) and negative (bottom row; WoF = −1) mood manipulations.

**Supplementary Figure 4.**
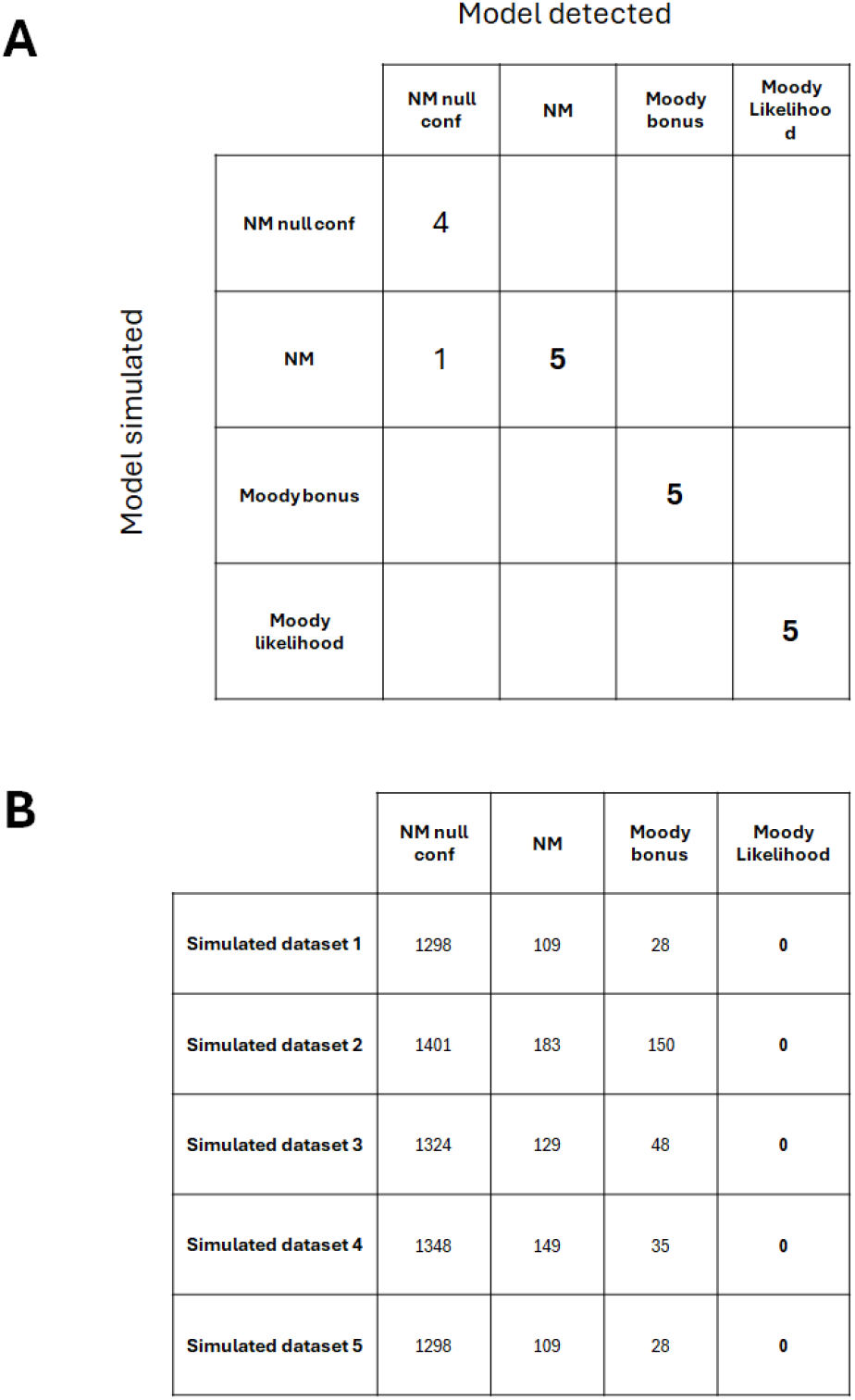
Validation of the model comparison procedure. We simulated 5 data sets using each model with parameter values from subjects’ real choices, adding Gaussian white noise, and then fit our models to each synthetic data set. **A)** Each cell shows how many of these synthetic datasets were detected as the model that generated the data. This analysis showed that the model comparison could always detect the model used for data simulation as the best-fitting model for our key models: non-moody accumulation (NM), moody bonus and Moody Accumulation models (5 times out of 5), although non-moody null confidence model was only detected 4 out of 5 times, it was only ever mis-detected as the non-moody model rather than one of the moody models. Importantly, the two moody models were never mis-detected, confirming specificity of the model comparison procedure. **B)** To further confirm the detectability of the winning Moody Accumulation model, we show the amount it was favoured in model detection analyses when it was indeed the generating model, compared to the other models. BIC scores here represent the model’s difference in BIC compared to the Moody Accumulation model (lower values signify higher model likelihood), within each of the 5 synthetic datasets.

**Supplementary Figure 5.**
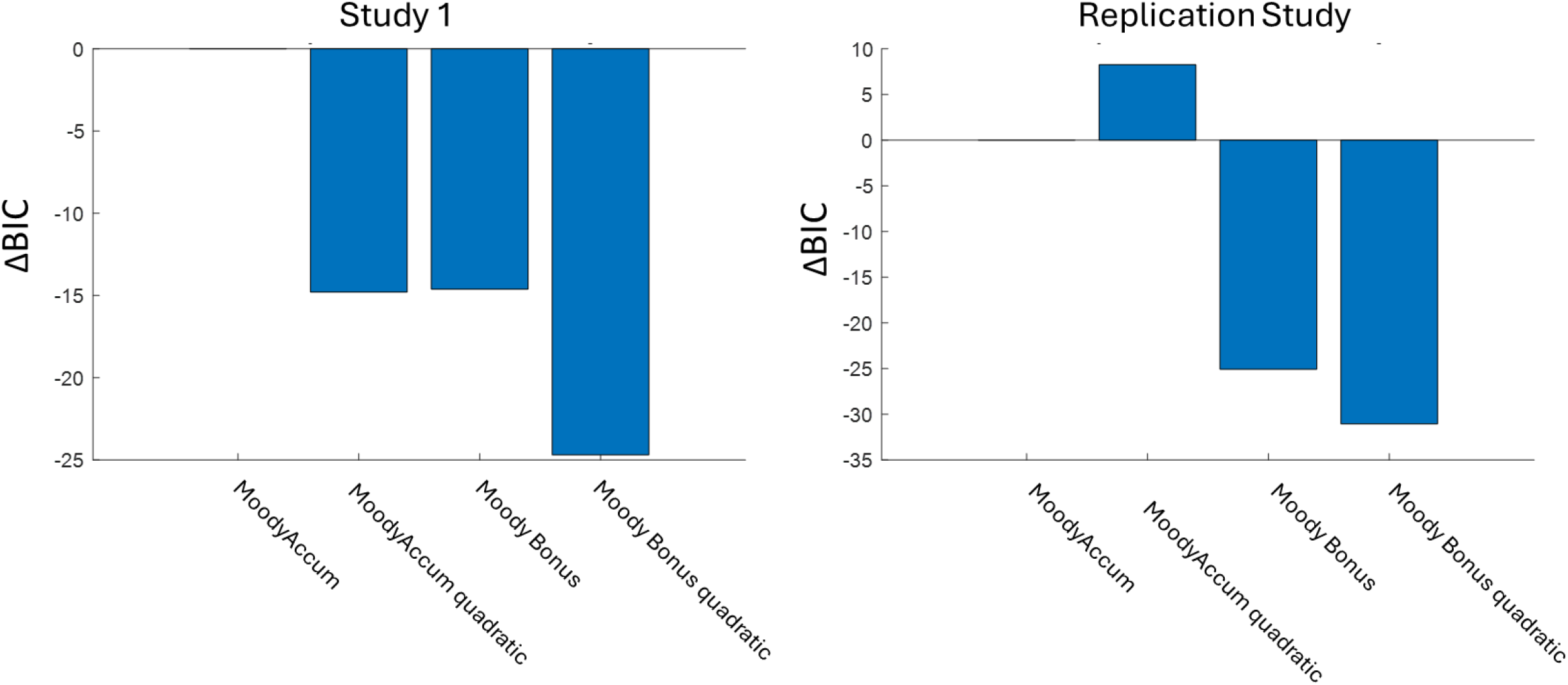
Model comparison of moody models with confidence specified as a linear versus quadratic function of subjective reward probability. Bars show ΔBIC relative to the linear *Moody Accumulation* model, defined such that positive values indicate improved model fit and negative values indicate worse fit (i.e., higher BIC). In Study 1, the quadratic variant Moody Accumulation model was less well supported (ΔBIC = −15), in the Replication study there was a small improvement (ΔBIC = 7).

